# Male age and *Wolbachia* dynamics: Investigating how fast and why bacterial densities and cytoplasmic incompatibility strengths vary

**DOI:** 10.1101/2021.06.01.446638

**Authors:** J. Dylan Shropshire, Emily Hamant, Brandon S. Cooper

**Author notes:** Correspondence to: J. Dylan Shropshire, Missoula, MT, 59801. **Author contributions:** JDS roles: Conceptualization, Data curation, Formal Analysis, Funding acquisition, Investigation, Methodology, Supervision, Validation, Visualization, Writing - original draft, Writing – review & editing. EH roles: Investigation, Validation, Writing – review & editing. BSC roles: Project administration, Conceptualization, Funding acquisition, Supervision, Writing – original draft, Writing – review & editing.

## Abstract

Endosymbionts can influence host reproduction and fitness to favor their maternal transmission. For example, endosymbiotic *Wolbachia* bacteria often cause cytoplasmic incompatibility (CI) that kills uninfected embryos fertilized by *Wolbachia*-modified sperm. Infected females can rescue CI, providing them a relative fitness advantage. *Wolbachia*-induced CI strength varies widely and tends to decrease as host males age. Since strong CI drives *Wolbachia* to high equilibrium frequencies, understanding how fast and why CI strength declines with male age is crucial to explaining age-dependent CI’s influence on *Wolbachia* prevalence. Here, we investigate if *Wolbachia* densities and/or CI gene (*cif*) expression covary with CI-strength variation and explore covariates of age-dependent *Wolbachia*-density variation in two classic CI systems. *w*Ri CI strength decreases slowly with *Drosophila simulans* male age (6%/ day), but *w*Mel CI strength decreases very rapidly (19%/ day), yielding statistically insignificant CI after only three days of *D. melanogaster* emergence. *Wolbachia* densities and *cif* expression in testes decrease as *w*Ri-infected males age, but both surprisingly increase as *w*Mel-infected males age, and CI strength declines. We then tested if phage lysis, Octomom copy number (which impacts *w*Mel density), or host immune expression covary with age-dependent *w*Mel densities—only host immune expression correlated with density. Together, our results identify how fast CI strength declines with male age in two model systems and reveal unique relationships between male age, *Wolbachia* densities, *cif* expression, and host immunity. We discuss new hypotheses about the basis of age-dependent CI strength and its contributions to *Wolbachia* prevalence.

**Importance:** *Wolbachia* are the most common animal-associated endosymbionts due in large part to their manipulation of host reproduction. Many *Wolbachia* cause cytoplasmic incompatibility (CI) that kills uninfected host eggs. Infected eggs are protected from CI, favoring *Wolbachia* spread in natural systems and in transinfected mosquito populations where vector-control groups use strong CI to maintain pathogen-blocking *Wolbachia* at high frequencies for biocontrol of arboviruses. CI strength varies considerably in nature and declines as males age for unknown reasons. Here, we determine that CI strength weakens at different rates with age in two model symbioses. *Wolbachia* density and CI gene expression covary with *w*Ri-induced CI strength in *Drosophila simulans*, but neither explain rapidly declining *w*Mel-induced CI in aging *D. melanogaster* males. Patterns of host immune gene expression suggest a candidate mechanism behind age-dependent *w*Mel densities. These findings inform how age-dependent CI may contribute to *Wolbachia* prevalence in natural systems and potentially in transinfected systems.

## Introduction

Reproductive parasites manipulate host reproduction to facilitate their spread in host populations. These endosymbiotic microbes may kill or feminize males or induce parthenogenesis to bias sex ratios favoring females (1). More frequently, reproductive parasites cause cytoplasmic incompatibility (CI) that reduces embryonic viability when aposymbiotic females mate with symbiont-bearing males (**Fig. 1A**) (2). Females harboring a closely related symbiont are compatible with CI-causing symbiotic males of the same strain, providing symbiont-bearing females a relative advantage that encourages symbiont spread to high frequencies in host populations (3–8). Divergent *Cardinium* (9), *Rickettsiella* (10), *Mesenet* (11), and *Wolbachia* (12) endosymbionts cause CI. *Wolbachia* are the most common, infecting 40-65% of arthropod species (13, 14). *Wolbachia* cause CI in at least ten arthropod orders (2), and pervasive CI directly contributes to *Wolbachia* spread and its status as one of the most common endosymbionts in nature.

**Figure 1.**
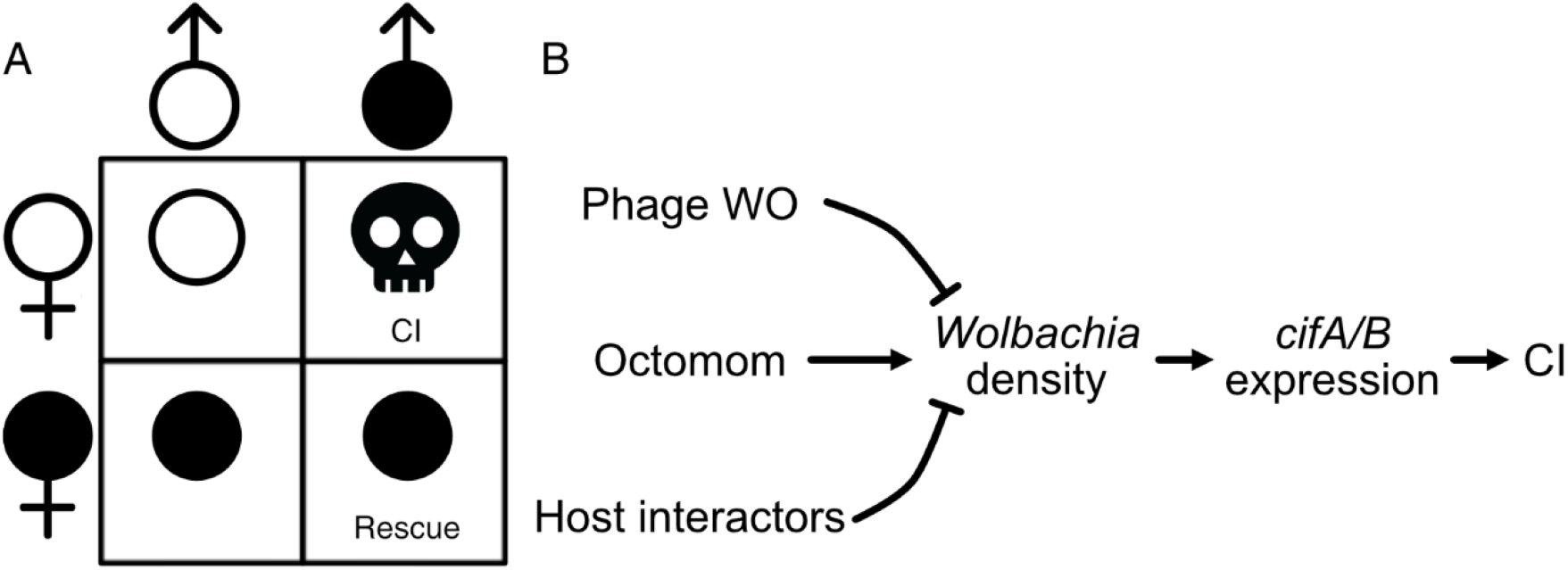
CI crossing relationships and potential causes of CI-strength variation. (A) CI causes embryonic death, measured as reduced embryo hatch when infected males (filled symbol) mate with uninfected females (unfilled symbol). All other crosses are compatible and have high embryonic hatching. Importantly, infected females maternally transmit *Wolbachia* and can rescue CI. (B) Schematic representation of factors that putatively impact *Wolbachia* densities, CI gene expression, and CI strength.

Within host populations, *Wolbachia* frequencies are governed by their effects on host fitness (15–20), maternal transmission efficiency (21–23), and CI strength (% embryonic death) (3, 5). CI strength varies from very weak to very strong and produces relatively low and high infection frequencies, respectively. For example, *w*Yak in *Drosophila yakuba* causes weak CI (∼15%) and tends to occur at intermediate and often variable frequencies (∼40-88%) in West Africa (22, 24). Conversely, *w*Ri in *D. simulans* causes strong CI (∼90%) and occurs at high and stable frequencies (e.g., ∼93% globally) (4, 25–27). In *D. melanogaster, w*Mel CI strength is relatively weak (28–30), contributing to considerably differing infection frequencies on multiple continents (31–35). In contrast, *w*Mel usually causes complete CI (no eggs hatch) in transinfected *Aedes aegypti* mosquitoes (36–39). Vector-control groups use strong CI induced by *w*Mel and other variants (e.g., *w*AlbB and *w*Pip) to either suppress mosquito populations through the release of infected males (40–45) or to drive pathogen-blocking *Wolbachia* to high and stable frequencies to inhibit pathogen spread (36, 46–49).

Despite CI’s importance for explaining *Wolbachia* prevalence in natural systems and reducing human disease transmission in transinfected mosquito systems, the mechanistic basis of CI-strength variation remains unresolved. Two hypotheses are plausible. First, the bacterial-density model predicts that CI is strong when bacterial density is high **(Fig. 1B)** (50). Indeed, *Wolbachia* densities positively covary with CI strength across *Drosophila*-*Wolbachia* associations (51, 52) and variable CI within strains (37, 38, 53–59). Second, the CI-gene-expression hypothesis predicts higher CI-gene expression contributes to stronger CI **(Fig. 1B)** (60). In *Drosophila*, two genes (*cifA* and *cifB*) associated with *Wolbachia*’s bacteriophage WO contribute to CI when expressed in testes (60–64), and one gene (*cifA*) rescues CI when expressed in ovaries (63–65). CI strength covaries with transgenic *cif* expression in *D. melanogaster* (60, 64), and natural *cif* expression covaries with CI strength in *Habrobracon* ectoparasitoid wasps (66). Bacterial density may explain CI strength via *cif* expression but may not perfectly align with CI strength since *Wolbachia* variably express *cifs* across conditions that impact CI strength (60). Thus, the bacterial-density and *cif*-expression hypotheses are not mutually exclusive. It remains unknown if *cif* expression is responsible for CI-strength variation and if it covaries with *Wolbachia* density in natural *Drosophila*-*Wolbachia* associations.

If symbiont density is a crucial factor governing CI strength, what governs the change in density? There are several plausible drivers of *Wolbachia*-density variation. First, phage WO is a temperate phage capable of cell lysis in some *Wolbachia* strains (66–70). Lytic phages form particles that burst through the bacterial cell membrane, killing the bacterial host. The phage density model proposes that as phage densities increase, *Wolbachia* densities decrease (**Fig. 1B**) (53). Temperature induced phage lysis covaries with lower *Wolbachia* densities and CI strength in some parasitoid wasps (53, 66), though it is unknown if phage lysis influences *Wolbachia* densities in any other systems. Second, *w*Mel *Wolbachia* have a unique ampliconic gene region composed of eight genes termed “Octomom” (71–75). Octomom copy number varies among *w*MelCS and *w*MelPop *Wolbachia* between host generations and positively covaries with *Wolbachia* densities **(Fig. 1B)**, but effects of Octomom-dependent *Wolbachia* densities on CI have not been investigated. Third, theory predicts that selection favors the evolution of host suppressors (6), as observed for male killing (76–79). Indeed, CI strength varies considerably across host backgrounds (24, 29, 39, 80–82), supporting a role for host genotype in CI-strength variation. The genetic underpinnings and mechanistic consequences of host suppression remain unknown, but two models have been proposed (2). The defensive model suggests that host CI targets diverge to prevent interaction with *cif* products, and the offensive model suggests that host products directly interfere with *Wolbachia* density or the proper expression of *cif* products (e.g., through immune regulation) **(Fig. 1B)**. Only a taxon-restricted gene of *Nasonia* wasps and host transcriptional activity in *Drosophila* have been functionally determined to contribute to *Wolbachia*-density variation (83, 84); thus, considerable work is necessary to uncover host determinants of variation in *Wolbachia*-density. Since *Wolbachia* densities significantly contribute to several phenotypes (54, 85), investigation of the causes of *Wolbachia*-density variation is sorely needed.

CI strength within *Wolbachia*-host systems covaries with several factors, including temperature (29, 37, 38, 53, 66), male mating rate (86, 87), male development time (88), rearing density (88), nutrition (89), paternal grandmother age (30), and male age (3, 18, 27, 29, 86). Male age does not always influence CI strength (90–92), but *w*Mel-infected *D. melanogaster* (29), *w*Ri-infected *D. simulans* (3, 18, 27), and other *Wolbachia*-infected hosts tend to cause weaker CI as males age (91, 93–95). CI seems to decline more slowly for *w*Ri (3, 18, 27) than for *w*Mel (3, 18, 27, 29), though the precise rates of CI-strength decline have not been estimated. While several factors might contribute to age-dependent CI strength, the mechanistic underpinnings of this phenotype remain unknown.

Here, we investigate rates of CI decline with male age and its mechanistic underpinnings in two classic *Wolbachia* CI systems: *w*Ri and *w*Mel (25, 28, 32). These *Wolbachia* diverged up to 6 million years ago and have unique *cif* repertoires (60, 63). We demonstrate that relative to *w*Ri, *w*Mel-induced CI strength declines more than three times faster, disappearing in a matter of days. We provide the first direct test of the *cif*-expression hypothesis in either system and the highest resolution investigation of *Wolbachia*-density variation across age to date. Our results suggest that *Wolbachia* density and *cif* expression in full-testes extracts cannot explain age-dependent CI-strength relationships across *Wolbachia*-host associations and motivate future work to investigate how host immunity could contribute to age-dependent *Wolbachia* densities. We discuss how these data inform our understanding of the causes of CI-strength variation, *Wolbachia*-density variation, and the consequences for *Wolbachia* prevalence in nature.

## Results

### How much does CI strength vary with age?

CI manifests as embryonic lethality (**Fig. 1A**). As such, we measured CI strength as the percent of embryos that hatch from a mating pair’s clutch of offspring—high compatibility corresponds with high hatching. Our experiments used males of different ages to test the impact of male age on CI strength. Here, we defined age as days since eclosion where males paired with females the day they eclosed were considered 0-days-old. For *w*Mel, we measured CI strength daily across the first three days of male age (**Fig. 2A**) and separately every two days across the first eight days of male age (**Fig. 2B**). This design enabled us to determine the rate of CI decline and the ages where males no longer cause significant CI. Crossing uninfected *D. melanogaster* females and males yielded high levels of compatibility (**Fig. 2A**; 95% confidence interval of the mean = 74 - 93%). Young 0-day-old *w*Mel-infected males induced strong CI when mated with uninfected females (95% interval = 9 - 27%). *w*Mel-infected females significantly rescued CI caused by infected 0-day-old males (95% interval = 87 - 92%, *P* = 1.74E-12).

**Figure 2.**
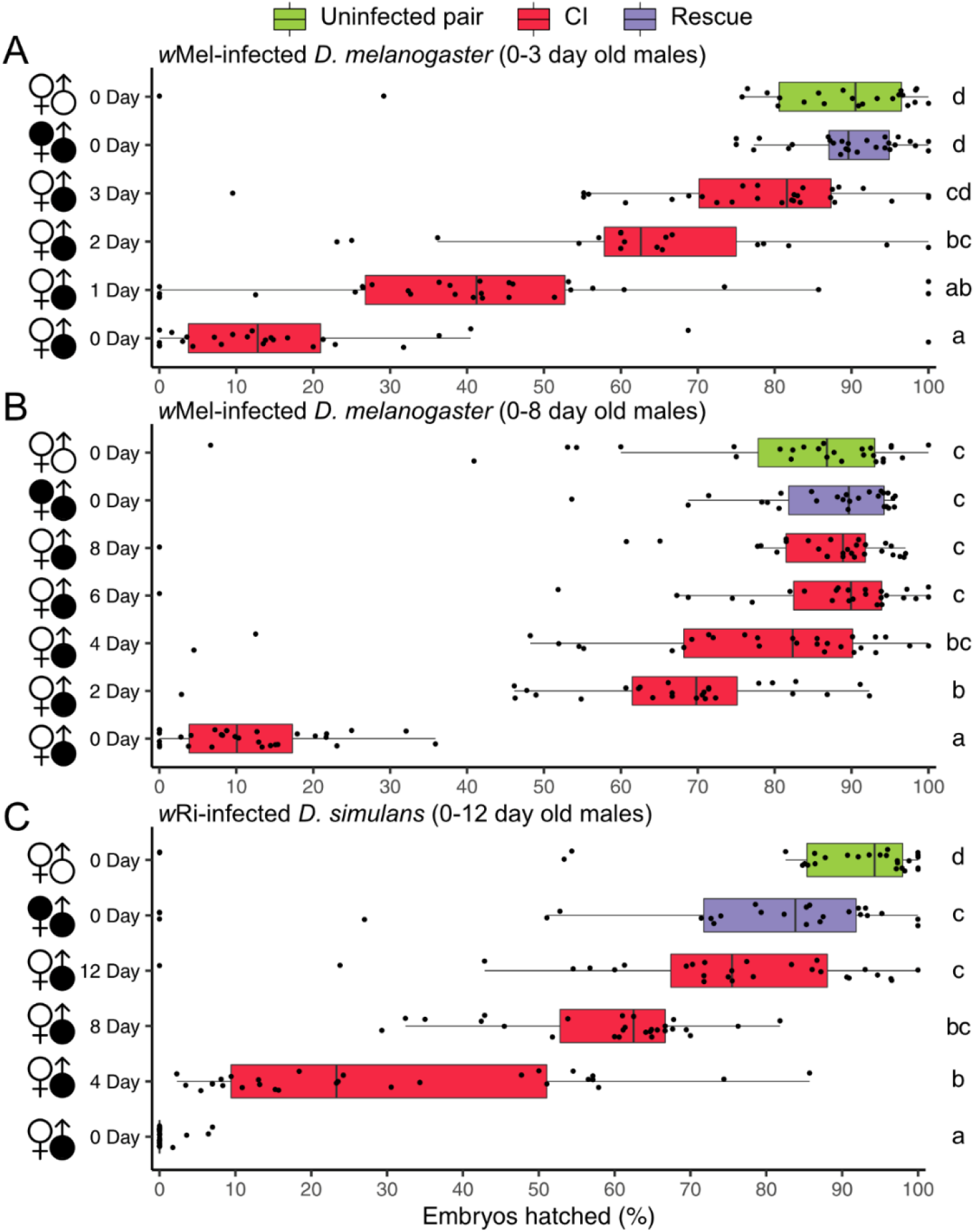
CI strength decreases as males age. (A) Hatch rate displaying CI strength with 0-, 1-, 2-, and 3-day-old *w*Mel-infected *D. melanogaster* males. (B) Hatch rate displaying CI strength with 0-, 2-, 4-, 6-, and 8-day-old *w*Mel-infected *D. melanogaster* males. (C) Hatch rate displaying CI strength with 0-, 4-, 8-, and 12-day-old *w*Ri-infected *D. simulans* males. Filled and unfilled sex symbols represent infected and uninfected flies, respectively. Male age is displayed to the right of the corresponding sex symbol. CI crosses are colored red, rescue crosses are purple, and uninfected crosses are green. Boxplots represent median and interquartile ranges. Letters to the right represent statistically significant differences based on α=0.05 calculated by Dunn’s test with correction for multiple comparisons between all groups —crosses that do not share a letter are significantly different. *P*-values are reported in **Table S1**. These data demonstrate that CI strength decreases with age in two *Wolbachia*-host associations and more slowly in *w*Ri-infected *D. simulans*.

Crosses using older 1- (95% interval = 31 - M51%), 2- (95% interval = 53 - 73%), and 3-day-old (95% interval = 69 - 83%) infected males trended toward progressively weaker CI (**Fig. 2A**). Average *w*Mel CI strength decreased daily by 19.3%: 22.8% from 0- to 1-day-old males, 21.8% from 1- to 2-day-old, and 13.4% from 2- to 3-day-old. Crosses between uninfected females and 3-day-old males (95% interval = 69 - 83%) did not cause significant CI, with egg hatch similar to the compatible uninfected (95% interval = 74 - 93%; *P* = 0.35) and rescue (95% interval = 87 - 92%; *P* = 0.19) crosses. These data highlight the rapid decline of *w*Mel CI strength with *D. melanogaster* male age.

In the experiment that includes older males (**Fig. 2B**), the uninfected cross also yielded high compatibility (95% interval = 72 - 88%). 0-day-old infected males caused strong CI when crossed with uninfected females (95% interval = 8 - 15%), and infected females significantly rescued 0-day-old CI (95% interval = 83 - 91%; *P* = 2.51E-12). Compatibility increased as males aged, where 2-day-old (95% interval = 59 - 73%) males caused significant CI and 4- (95% interval = 66 - 83%), 6- (95% interval = 76 - 92%), and 8-day-old (95% interval = 77 - 91%) infected males did not significantly inhibit egg hatch relative to the compatible uninfected cross (*P* = 1 in all cases) (**Fig 2B**). Average *w*Mel CI strength decrease by approximately 19.3% each day as *D. melanogaster* males aged, but this rate of decrease slowed each day, such that CI was no longer statistically detectable once males were 3-days-old.

Next, we assessed age-dependent CI in *w*Ri-infected *D. simulans* (**Fig. 2C**). As expected, uninfected *D. simulans* females and males were compatible (95% interval = 74 - 94%). Young 0-day-old *w*Ri-infected males caused strong CI when mated with uninfected females (95% interval = 0 - 1%), and infected females significantly rescued 0-day-old CI (95% interval = 59 - 84%; *P* = 1.83E-10). Older 4- (95% interval = 21 - 39%), 8- (95% interval = 54 - 64%), and 12-day-old (95% interval = 64 - 82%) infected males induced progressively weaker CI as males aged. Average *w*Ri CI strength decreased by about 6.0% per day: 29.1% (7.3%/ day) from 0-day-old to 4-day-old males, 29.0% (7.3%/ day) from 4-day-old to 8-day-old, and 14.0% (3.5%/ day) from 8-day-old to 12-day-old. These data support a strong effect of *D. simulans* male age on *w*Ri CI strength, but the daily decrease is more than three times slower than what we observed for *w*Mel CI strength decline as *D. melanogaster* males age.

### What causes CI strength to vary with age?

The bacterial-density and CI-gene-expression hypotheses are both proposed to explain CI-strength variation. These hypotheses predict that *Wolbachia* density and/or *cif* expression positively covary with CI strength. To elucidate the causes of declining CI strength with male age, we tested both hypotheses in the context of rapidly declining *w*Mel CI strength and more slowly declining *w*Ri CI strength.

#### Bacterial density differentially covaries with age between species

We tested the bacterial density hypothesis by dissecting testes from siblings of flies used in our CI assays above, extracting DNA, and measuring the relative abundance of a single-copy *Wolbachia* gene (*ftsZ*) relative to a single-copy ultraconserved element (UCE) (96) of *Drosophila* via qPCR. We selected a random infected sample from the 0-day-old age group as the reference for all fold change analyses within each experiment. We report all qPCR data as fold change relative to this control. Surprisingly, 0-day-old *D. melanogaster* testes had low *w*Mel density (**Fig. 3A**; 95% interval = 0.53 - 1.01 fold change), and older 2- (95% interval = 0.92 - 1.11), 4- (95% interval = 0.96 - 1.72), 6- (95% interval = 1.17 - 1.49), and 8-day-old (95% interval = 1.19 - 1.51) infected testes had progressively higher *w*Mel densities (**Fig. 3A**). *w*Mel densities were significantly different among age groups according to a Kruskal-Wallis test (**Fig. 3A**; *P* = 1.1E-03). To test for a correlation between *w*Mel densities and CI strength, we performed Pearson (r_p_) and Spearman (r_s_) correlations on the relationship between *w*Mel fold change against median hatch rates from the associated age groups. *w*Mel densities were significantly positively correlated with increasing compatibility (**Table S3**; r_p_ = 0.75, *P* = 5.5E-06; r_s_ = 0.77, *P* = 2.3E-06). *w*Mel densities also covaried with age (**Fig. S1A**; *P* = 0.02) and correlated with increasing compatibility (**Table S3**; r_p_ = 0.64, *P* = 7.7E-04; r_s_ = 0.64, *P* = 7.4E-04) in the younger 0-, 1-, 2-, and 3-day-old *D. melanogaster* age group. This result was contrary to our prediction that higher *w*Mel densities would be correlated with stronger CI and lower compatibility.

**Figure 3.**
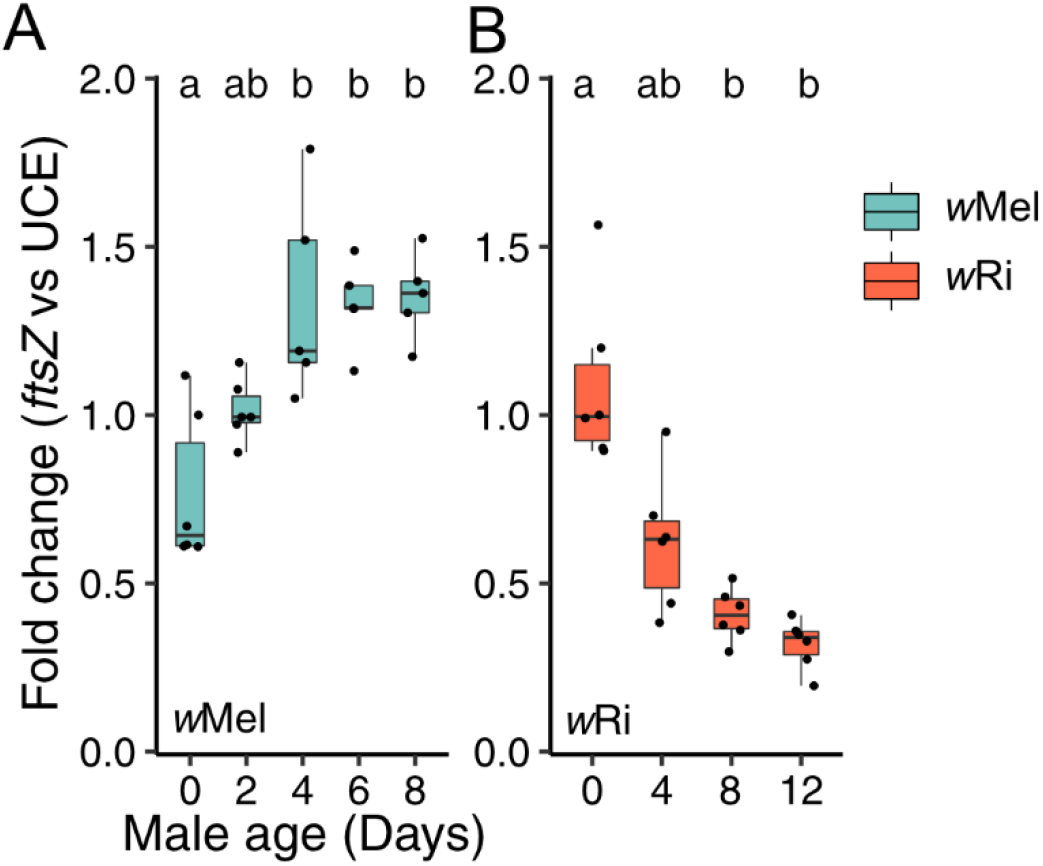
Testing the bacterial density model for CI-strength variation. Fold change in testes across male age for the relative expression of (A) *w*Mel *ftsZ* to *D. melanogaster* UCE and (B) *w*Ri *ftsZ* to *D. simulans* UCE. Letters above data represent statistically significant differences based on α=0.05 calculated by a Dunn’s test with correction for multiple comparisons between all groups—crosses that do not share a letter are significantly different. Fold change was calculated as 2^-ΔΔcq^. We selected a random infected sample from the youngest 0-day-old age group as the reference for all fold change analyses within each experiment. *P*-values are reported in **Table S1**. These data demonstrate that *Wolbachia* density differentially covaries with age between *Wolbachia*-host associations.

Next, we tested the bacterial density model in *w*Ri-infected *D. simulans*. In contrast to *w*Mel, *w*Ri-infected 0-day-old (95% interval = 0.82 - 1.36) *D. simulans* testes had the highest *w*Ri densities and they consistently decreased in 4- (95% interval = 0.41 - 0.83), 8- (95% interval 0.41 - 0.83), and 12-day-old (95% interval = 0.24 - 0.40) testes (**Fig. 3B**). *w*Ri densities were significantly different among *D. simulans* age groups (*P* = 3.9E-04) and were significantly negatively correlated with increasing compatibility (**Table S3**; r_p_ = -0.84, *P* = 2.4E-07; r_s_ = -0.89, *P* = 6.9E-09).

In conclusion, these data fail to support the bacterial-density hypothesis for age-dependent CI-strength variation in *w*Mel-infected *D. melanogaster* but support the hypothesis in *w*Ri-infected *D. simulans*. Thus, *Wolbachia* densities from full-testis extracts cannot explain age-dependent CI across *Wolbachia*-host associations, suggesting that other factors contribute to these patterns.

#### cif expression varies with age, but the direction differs between strains

*cif* expression is hypothesized to control CI-strength variation within *Wolbachia*-host associations (2, 60). *cif* loci are classified into five different phylogenetic clades called ‘Types’ (60, 97–99). *w*Mel has a single pair of Type I *cifs*, and *w*Ri has two identical pairs closely related to the *w*Mel copy plus a divergent Type 2 pair (60). We investigated three questions regarding *cif* expression. First, does *cif* expression change relative to the host as males age? We expected *cif* expression per host cell would be the key determinant of CI-strength variation. To test this, we used RT-qPCR to measure the transcript abundance of *cifA* and *cifB* and compared their expression to β Spectrin (*βspec*), a *Drosophila* membrane protein with invariable expression with age (see Materials and Methods for details). Second, does *cif* expression decrease relative to *Wolbachia* as males age? Since *w*Mel densities increase with male age, *w*Mel would need to express *cif*_*wMel[T1]*_ at lower levels in older males to allow *cif*_*wMel[T1]*_ to decrease relative to the host. Finally, does *cifA* expression change relative to *cifB* as males age? Evidence of differential localization of *cif* loci that covaries with age might indicate more complex determinants of age-dependent CI based on the relative abundance of these products.

We started by investigating these question in *w*Mel-infected *D. melanogaster*. Contrary to our first prediction, the relative expression of *cifA*_*wMel[T1]*_ to *D. melanogaster βspec* was lowest in 0-day-old infected males (95% = 1.1 - 1.6) and consistently increased in 2- (95% interval = 1.5 - 3.2), 4- (95% interval = 1.9 - 2.3), 6- (95% interval = 2.1 - 2.8), and 8-day-old (95% interval = 0.9 - 3.8) testes (**Fig. 4A**). Relative expression of *cifA*_*wMel[T1]*_ to *βspec* significantly varied across male age (*P* = 8.4E-03) and was significantly positively correlated with increasing compatibility (**Table S3**; r_p_ = 0.61, *P* = 6.4E-04; r_s_ = 0.59, *P* = 9.7E-04). Comparably, relative expression of *cifB*_*wMel[T1]*_ to *βspec* significantly increased with male age (**Fig. S2A**; *P* = 7.3E-03). Analysis of raw quantification cycle (C_q_) variation with age supports increased *cifA*_*wMel[T1]*_ (**Fig. S2C**; *P* = 3.1E-04) and *cifB*_*wMel[T1]*_ (**Fig. S2D**; *P* = 1.1E-03) expression; *βspec* C_q_ does not vary with age (**Fig. S2E**; *P* = 0.1) and *ftsZ* C_q_ significantly decreases with age (**Fig. S2F**; *P* = 1.3E-04). Thus, we report for the first time that *cif*_*wMel[T1]*_ expression relative to the host in full-testes extracts is not sufficient to explain CI-strength variation, leading us to reject the hypothesis that *cif*_*wMel[T1]*_ expression in full-testes extracts can explain age-dependent *w*Mel CI strength.

**Figure 4.**
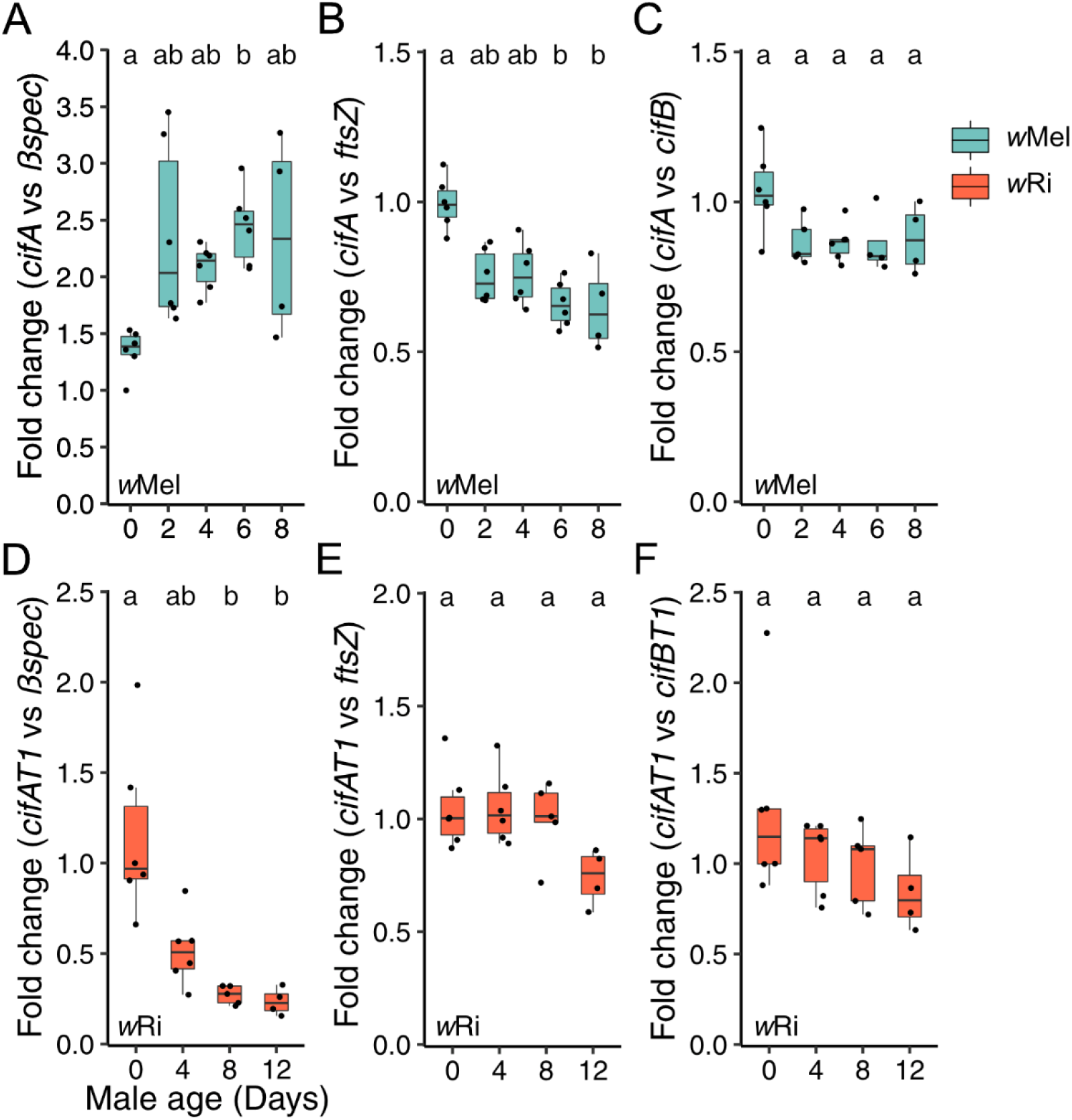
Testing the *cif*-expression hypothesis for CI-strength variation. Fold change in testes across male age for the relative expression of (A) *cifA*_*wMel[T1]*_ to *D. melanogaster βspec*, (B) *cifA*_*wMel[T1]*_ to *w*Mel *ftsZ*, (C) *cifA*_*wMel[T1]*_ to *cifB*_*wMel[T1]*_, (D) *cifA*_*wRi[T1]*_ to *D. simulans βspec*, (E) *cifA*_*wRi[T1]*_ to *w*Ri *ftsZ*, and (F) *cifA*_*wRi[T1]*_ to *cifB*_*wRi[T1]*_. Letters above data represent statistically significant differences based on α=0.05 calculated by Dunn’s test with correction for multiple comparisons between all groups —crosses that do not share a letter are significantly different. Fold change was calculated as 2^-ΔΔcq^. We selected a random infected sample from the youngest 0-day-old age group as the reference for all fold change analyses within each experiment. *P*-values are reported in **Table S1**. These data demonstrate that age-dependent *cif* expression is variably related to host expression, *cif*_*wMel[T1]*_ expression decreases per *Wolbachia* with age, and *cifA/B* relative expression only marginally decreases with age in both systems.

Next, we investigated our second question: does *cif*_*wMel[T1]*_ expression vary relative to *Wolbachia* as males age? Indeed, relative expression of *cifA*_*wMel[T1]*_ to *w*Mel *ftsZ* was highest in 0-day-old infected-*D. melanogaster* testes (95% interval = 0.9 - 1.1), and consistently decreased in 2- (95% interval = 0.7 - 0.8), 4- (95% interval = 0.7 - 0.9), 6- (95% interval = 0.6 - 0.7), and 8-day-old (95% interval = 0.4 - 0.9) testes (**Fig. 4B**). Relative expression of *cifA*_*wMel[T1]*_ to *w*Mel *ftsZ* significantly varied with age (*P* = 2.9E-03) and was significantly correlated with increasing compatibility (**Table S3**; r_p_ = -0.8, *P* = 4.0E-07; r_s_ = -0.7, *P* = 3.5E-05). Relative expression of *cifB*_*wMel[T1]*_ to *w*Mel *ftsZ* did not significantly covary with age (**Fig. S2B**; P = 0.3), but was significantly correlated with increasing compatibility (**Table S3**; r_p_ = -0.42, *P* = 3.7E-02; r_s_ = - 0.46, *P* = 2.2E-02). In summary, *cif*_*wMel[T1]*_ expression decreased relative to a *Wolbachia* housekeeping gene with age, consistent with prior reports that *w*Mel expression of *cifA*_*wMel[T1]*_ and *cifB*_*wMel[T1]*_ decrease as males age (60). However, since *cif*_*wMel[T1]*_ expression did not decrease relative to the host with age, we conclude that the decrease in *cif*_*wMel[T1]*_ expression per *Wolbachia* is insufficient to overcome the increase in *cif*_*wMel[T1]*_ expression caused by increased *w*Mel density in full-testes extracts.

Finally, we tested if the relative expression of *cifA*_*wMel[T1]*_ to *cifB*_*wMel[T1]*_ varied with age. Intriguingly, *cifA*/*B*_*wMel[T1]*_ relative expression did not significantly covary with age (**Fig. 4C**; *P* = 0.09), but was positively correlated with decreasing CI strength (**Table S3**; r_p_ = -0.61, *P* = 1.3E-03; r_s_ = -0.46, *P* = 0.021). In summary, these data suggest that *cif*_*wMel[T1]*_ expression per *w*Mel decreases as males age, that *cifA*_*wMel[T1]*_ expression decreases marginally faster than *cifB*_*wMel[T1]*_, and that overall *cif*_*wMel[T1]*_ expression increases relative to the host as males age and CI strength decreases. This is the first report that CI strength is decoupled from *Wolbachia* densities and *cif* expression in testes.

Next, we investigated the *cif*-expression hypotheses in *w*Ri. We predicted that *cif*_*wRi[T1]*_ and/or *cif*_*wRi[T2]*_ expression would decrease relative to host expression. Since *w*Ri density decreased with age, *cif* expression per *w*Ri would not need to change to accomplish this shift in relative expression. As predicted, relative expression of *cifA*_*wRi[T1]*_ to *D. simulans βspec* was highest in infected 0-day-old (95% interval = 0.7 - 1.7) testes, and declined in 4- (95% interval = 0.1 - 0.4), 8- (95% interval = 0.3 - 0.7), and 12-day-old (95% interval = 0.2 - 0.3) testes (**Fig. 4D**). Relative expression of *cifA*_*wRi[T1]*_ to *D. simulans βspec* significantly covaried with age (*P* = 1.2E-03) and was significantly correlated with decreasing CI strength (**Table S3**; r_p_ = -0.76; r_s_ = - 0.88). Similarly, relative expression of *cifB*_*wRi[T1]*_ (**Fig. S3A**; *P* = 2.3E-03), *cifA*_*wRi[T2]*_ (**Fig. S3C**; *P* = 1.9E-03), and *cifB*_*wRi[T2]*_ (**Fig. S3E**; *P* = 1.2E-03) to *D. simulans βspec* also decreased with age and each were significantly correlated with decreasing CI strength (**Table S3**). These results support the *cif*-expression hypothesis for age-dependent CI in *w*Ri.

As with *w*Mel-infected *D. melanogaster* testes, relative expression of *cifA*_*wRi[T1]*_ to *w*Ri *ftsZ* significantly covaried with male age (**Fig. 4E**; *P* = 4.1E-02) and was significantly correlated with decreasing CI strength (**Table S3**; r_p_ = -0.47, *P* = 0.032; r_s_ = -0.47, *P* = 0.033). However, 0- (95% interval = 0.9 - 1.2), 4- (95% interval = 0.9 - 1.2), and 8-day-old (95% interval = 0.8 - 1.2) testes had similar expression patterns, suggesting that expression in 12-day-old (95% interval = 0.5 - 0.9) testes drove this significant difference; though, a Dunn’s test was unable to identify significantly different pairs (**Fig. 4E**). Conversely, *cifB*_*wRi[T1]*_ (**Fig. S3B**; *P* = 0.6), *cifA*_*wRi[T2]*_ (**Fig. S3D**; *P* = 0.2), and *cifB*_*wRi[T2]*_ (**Fig. S3F**; *P* = 0.2) expression relative to *w*Ri *ftsZ* did not vary with age or decreasing CI strength (**Table S3**).

Finally, as with *w*Mel, we investigated the relationship between *cifA* and *cifB* expression in *w*Ri across age and found similar results where *cifA*_*wRi[T1]*_ expression relative to *cifB*_*wRi[T1]*_ expression did not significantly vary with male age (**Fig. 4F**; *P* = 0.2) but did significantly correlate with increasing compatibility (**Table S3**; r_p_ = -0.44, *P* = 0.045; r_s_ = -0.46, *P* = 0.035). Relative expression of *cifA*_*wRi[T1]*_ to *cifA*_*wRi[T2]*_ expression did not covary with age (**Fig. S3G**; *P* = 0.6) or increasing compatibility (**Table S3**; r_p_ = 0.01, *P* = 0.96; r_s_ = -0.05, *P* = 0.84). Analysis of raw C_q_ values supported decreasing *cifA*_*wRi[T1]*_ (**Fig. S3H**; *P* = 1.0E-03), *cifB*_*wRi[T1]*_ (**Fig. S3I**; *P* = 8.1E-04), *cifA*_*wRi[T2]*_ (**Fig. S3J**; *P* = 1.8E-03), and *cifB*_*wRi[T2]*_ (**Fig. S3K**; *P* = 1.7E-03) expression with male age; *D. simulans βspec* C_q_ did not vary with age (**Fig. S3L**; *P* = 0.6) and *w*Ri *ftsZ* C_q_ significantly increased with age (**Fig. S3M**; *P* = 8.9E-04). In summary, *cif*_*wRi*_ expression significantly decreased with age in *w*Ri testes, *cifA*_*wRi[T1]*_ expression decreased marginally faster than *cifB*_*wRi[T1]*_ expression, and there was a small decrease in *cifA*_*wRi[T1]*_ expression relative to *w*Ri but other *cif*_*wRi*_ loci do not follow similar trends.

In conclusion, we found that *w*Mel *cif* expression did not explain age-dependent CI-strength variation. More specifically, *w*Mel’s expression of *cif* genes decreased with age (60), relative *w*Mel and *w*Ri *cifA*-to-*cifB* expression varied marginally with age, and *cif* expression dynamics varied considerably across male age and differed between *w*Mel- and *w*Ri-infected hosts.

### What causes Wolbachia density to vary with age?

We found that *Wolbachia* densities from full-testes extracts significantly increased with male age in *w*Mel-infected *D. melanogaster* and significantly decreased with male age in *w*Ri-infected *D. simulans*. The causes of age-dependent *Wolbachia*-density variation have not been explored. We tested three hypotheses. Namely, that phage lytic activity, Octomom copy number, or host immune expression may govern age-dependent *Wolbachia* densities.

#### Phage density does not covary with age-dependent Wolbachia density

The phage density hypothesis predicts that *Wolbachia* density negatively covaries with phage lytic activity (53). Since phage lysis corresponds with increased phage copy number (53, 66), we tested the phage density model by measuring the relative abundance of phage to *Wolbachia ftsZ* using qPCR. *w*Mel and *w*Ri each harbor a unique set of phage haplotypes: *w*Mel has two phages (WOMelA and WOMelB), and *w*Ri has four (WORiA-C, WORiB is duplicated) (100). At least one of *w*Mel’s phages is capable of particle production, but it is unknown if *w*Ri’s phages yield viral particles (70). We monitored WOMelA and WOMelB of *w*Mel simultaneously using primers that target homologs present in a single copy in each phage. Conversely, we monitored WORiA, WORiB, and WORiC separately since shared homologs are too diverged to make suitable qPCR primers that match multiple phage haplotypes.

First, we evaluated the phage density model for *w*Mel. We predicted the relative abundance of WOMelA/B to decrease with *D. melanogaster* male age since *w*Mel density increases with age. However, there was no change in WOMelA/B abundance relative to *w*Mel *ftsZ* as males aged (**Fig. 5A**; *P* = 0.3), while WOMelA/B abundance relative to *D. melanogaster* UCE increased similar to *w*Mel density (**Fig. S4A**; *P* = 3.0E-04). Relative phage abundance was not significantly correlated with increasing compatibility (**Table S3**; r_p_ = -0.065, *P* = 0.75; r_s_ = 0.17, *P* = 0.39). Similarly, WOMelA/B significantly varied with age relative to UCE (**Fig. S4B**; *P* = 0.049) but not *w*Mel *ftsZ* (**Fig. S4C**; *P* = 0.15) in the 0-, 1-, 2-, and 3-day-old age experiment.

**Figure 5.**
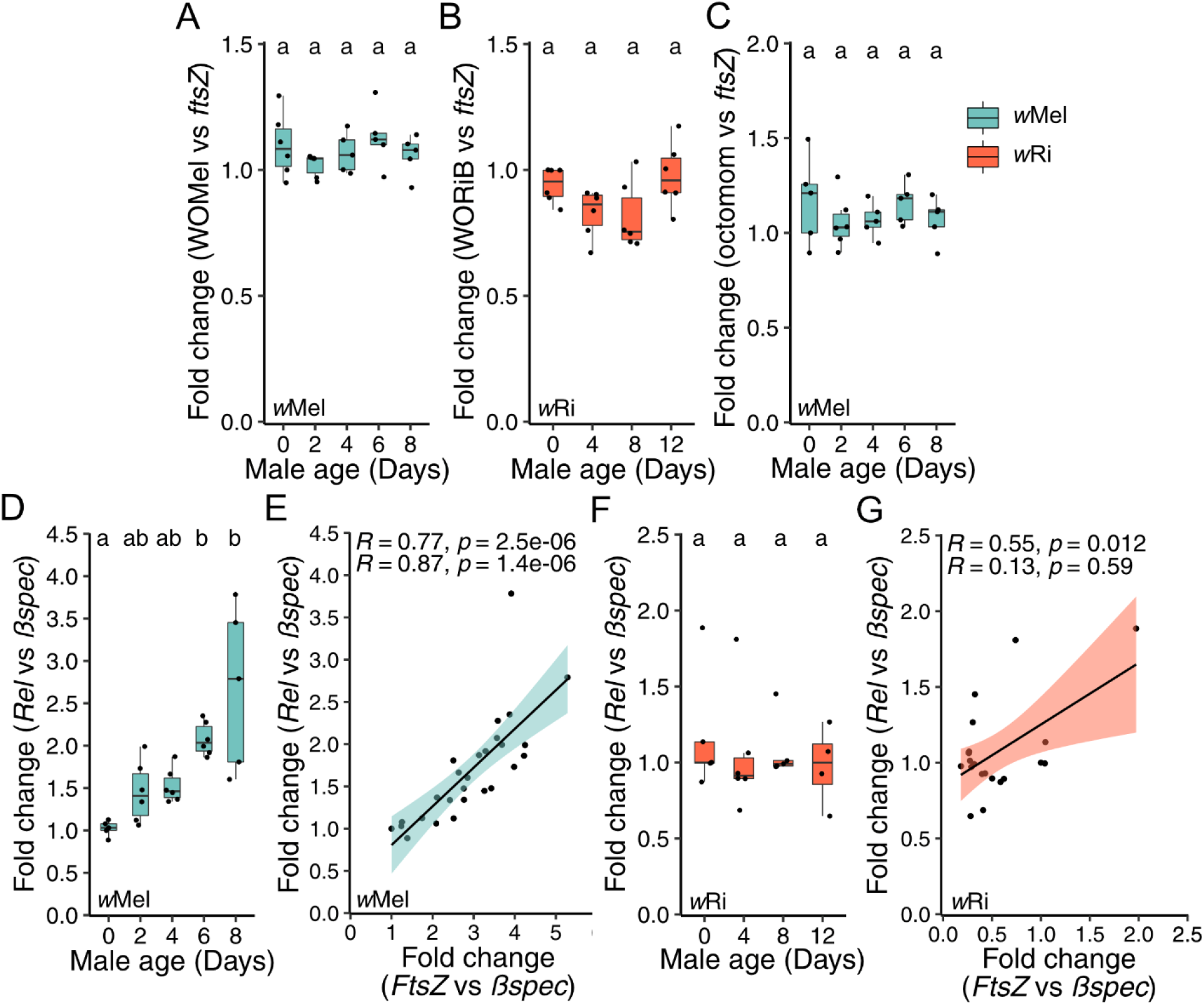
Testing the phage density, Octomom, and host immunity hypotheses for age-dependent *Wolbachia*-density variation. Fold change in testes across male age for the relative abundance or expression of (A) WOMelA/B to *w*Mel *ftsZ*, (B) WORiB to *w*Ri *ftsZ*, (C) Octomom gene WD0509 to *w*Mel *ftsZ*, (D) *D. melanogaster Rel* to *Bspec*, and (F) *D. simulans Rel* to *Bspec*. Correlation between the relative expression of *Rel* to *Bspec* and *ftsZ* to *Bspec* for (E) *w*Mel and (F) *w*Ri. Letters above data represent statistically significant differences based on α=0.05 calculated by Dunn’s test with correction for multiple comparisons between all groups — crosses that do not share a letter are significantly different. (E, G) Pearson (top) and Spearman (bottom) correlations are reported. Linear regressions are plotted with 95% confidence intervals. Fold change was calculated as 2^-ΔΔcq^. We selected a random infected sample from the youngest 0-day-old age group as the reference for all fold change analyses within each experiment. *P*-values are reported in **Table S1**. These data demonstrate that age-dependent *Wolbachia* densities are not controlled by phage WO lysis or Octomom copy number, but are correlated with *Rel* expression in *D. melanogaster* and less so in *D. simulans*.

Next, we predicted that WORi phage abundance would increase with decreasing *w*Ri densities across *D. simulans* male age if governed by the phage density model. As with *w*Mel in *D. melanogaster*, relative WORiB to *w*Ri *ftsZ* abundance did not significantly covary with male age (**Fig. 5B**; *P* = 0.053) or correlate with increasing compatibility (**Table S3**; r_p_ = 0.032, *P* = 0.88; r_s_ = 0.12, *P* = 0.58). Relative WORiB to *D. simulans* UCE abundance increased with age, similar to *w*Ri density (**Fig. S4D**; *P* = 4.4E-04). Comparably, WORiA (**Fig. S4E**; *P* = 0.3) and WORiC (**Fig. S4F**; *P* = 0.4) abundance relative to *w*Ri did not vary with male age. These data suggest that phage WO is unrelated to age-dependent *Wolbachia*-density variation in *w*Mel and *w*Ri.

#### Octomom does not vary with age-dependent wMel density

Only very closely related *w*Mel variants encode all eight Octomom genes (e.g., *w*Mel, *w*MelCS, *w*MelPop). The relative abundance of Octomom to *Wolbachia* genes positively covaries with *w*MelCS and *w*MelPop density (71–75), commonly changing between host generations. A pair of repeat regions flank the Octomom genes and are hypothesized to be involved in Octomom amplification. In *w*Mel, the 3’ repeat region has a transposon insertion that likely prevents Octomom amplification (71). As such, we predicted that Octomom copy number would be invariable with age. We tested if Octomom copy number variation correlated with age-dependent *w*Mel density variation using qPCR. Indeed, the relative abundance of an Octomom gene (WD0509) to *w*Mel *ftsZ* did not covary with male age (**Fig. 5C**; *P* = 0.53) or correlate with increasing compatibility (**Table S3**; r_s_ = -0.19, *P* = 0.36; r_s_ = 0.1, P = 0.61). Similar results were observed in 0-, 1-, 2-, and 3-day-old *w*Mel-infected males **(Fig. S1B; Table S3)**. We conclude that Octomom copy number is unrelated to the age-dependent increase in *w*Mel densities.

#### Relish expression is positively correlated with age-dependent wMel, but not wRi, densities

Theory predicts that natural selection favors the evolution of host genes that suppress CI (6). Manipulation of *Wolbachia* densities is one mechanism that may drive CI suppression (2). Since the immune system is designed to control bacterial loads, we investigated the role of the host immune system in *Wolbachia*-density variation across male age. The immune deficiency (Imd) pathway is broadly involved in defense against gram-negative bacteria like *Wolbachia* (101). Bacteria activate the Imd pathway by interacting with peptidoglycan (PG) recognition proteins which start a signal cascade that results in the expression of the NF-κB transcription factor Relish (*Rel*). Relish then activates antimicrobial peptide production. *Wolbachia* lack the full suite of genes needed to synthesize PG (102–104) but can express the PG precursor lipid II which is sufficient to activate the Imd pathway (104, 105).

We predicted that *D. melanogaster* Relish expression and *w*Mel density would be correlated if the Imd pathway is involved in *w*Mel density regulation. Indeed, relative expression of Relish to *βspec* significantly varied among age groups (*P* = 6.1E-4). However, relative expression of Relish to *βspec* was lowest in 0-day-old (95% interval = 0.9 - 1.1) infected testes and consistently increases in 2- (95% interval = 1.1 - 1.8), 4- (95% interval = 1.3 - 1.7), 6- (95% interval = 1.9 - 2.3), and 8-day-old (95% interval = 1.5 - 3.9) testes (**Fig. 5D**). Relish expression was significantly positively correlated with *w*Mel density within testes samples (**Fig. 5E**; r_p_ = 0.77, *P* = 2.5E-06; r_s_ = 0.87, *P* = 1.4E-06). In summary, *w*Mel density was strongly correlated with increasing Relish expression, directly contrary to our prediction.

Conversely, relative expression of *D. simulans* Relish to *βspec* did not significantly covary with age (**Fig. 5F**; *P* = 0.7), but remained positively correlated with the relative expression of *w*Ri *ftsZ* to *βspec* within testes samples according to Pearson, but not Spearman, analyses (**Fig. 5G**; r_p_ = 0.55, *P* = 0.012; r_s_ = 0.13, *P* = 0.59). In summary, Relish expression is positively correlated with age-dependent *w*Mel densities in *D. melanogaster*, but less so in *w*Ri-infected *D. simulans*, supporting a role for the Imd pathway in the regulation of at least *w*Mel density variation. However, more work is necessary to determine if the correlation between age-dependent immune expression and *Wolbachia* density in testes is causatively associated.

## Discussion

Within *Wolbachia*-host systems, several factors influence CI strength (29, 30, 37, 38, 53, 66, 86–89), but male age can be particularly impactful (3, 18, 27, 29). Our results determine how fast and investigate why CI strength declines as males age. First, we estimate that CI-strength decreases rapidly for *w*Mel-infected *D. melanogaster* (19%/ day), becoming statistically insignificant when males reach three days old. In contrast, *w*Ri causes intense CI that declines more slowly (6%/ day), resulting in statistically significant CI through at least the first 12 days of *D. simulans* male life. Second, *Wolbachia* densities and *cif* expression from full-testes extracts increase in *w*Mel-infected *D. melanogaster* and decrease in *w*Ri-infected *D. simulans* as males age and CI weakens. These results indicate that bacterial density and CI gene expression in full-testes extracts cannot fully account for age-dependent CI strength across host-*Wolbachia* associations. Third, while WO phage activity and Octomom copy number cannot explain *Wolbachia*-density variation, *D. melanogaster* immune expression covaries with *w*Mel densities, suggesting the host immune system may contribute to age-dependent *Wolbachia* density in *D. melanogaster*, but much less so in *D. simulans*. Notably, transcript-based data (e.g., *cif* and Relish) described here are subject to the caveat that mRNA levels may not correlate perfectly with protein expression or activity. Future proteomics analyses will be needed to confirm these trends hold at the protein level. We discuss how our discoveries inform the basis of age-dependent CI-strength variation, how multiple mechanistic underpinnings may govern age-dependent *Wolbachia* densities, and how age-dependent CI may contribute to *Wolbachia* frequency variation observed in nature.

### Wolbachia density and CI-gene expression in full-testes extracts do not fully explain age-dependent CI-strength variation

Despite support that CI strength is linked to *Wolbachia* density and *cif* expression across and within systems (37, 38, 51–54, 60, 66), our observations add to a growing body of literature suggesting *Wolbachia* densities in adult testes (30, 88) and, for the first time, *cif* expression, are insufficient to explain CI-strength variation broadly. We discuss three hypotheses to explain the disconnect between *Wolbachia* density and *cif* expression in full-testes extracts and CI strength with male age. Note, however, that these results may also be explained by a decoupling of *cif* transcription and protein translation, which will require future proteomics analyses to investigate.

First, the localization and density of *Wolbachia* and *cif* products within specific cells in testes may more accurately predict CI strength. Indeed, the proportion of infected spermatocyte cysts covaries with CI strength in natural and transinfected combinations of CI-inducing *Wolbachia* and *D. melanogaster, D. simulans, D. yakuba, D. teissieri*, and *D. santomea* (51, 52). Intriguingly, two *w*Ri-infected *D. simulans* strains whose *Wolbachia* cause variable CI did not have different *Wolbachia* densities according to qPCR, but the number of infected sperm cysts covaried with CI between strains (106). Thus, *Wolbachia* densities in full-testes extracts may not reflect the cyst infection frequency, but it is unknown how generalizable this discrepancy is across or within *Wolbachia*-host associations with variable CI strengths. It seems plausible that while *w*Mel densities increase in the testes as males age, the proportion of infected spermatocytes could decrease. Notably, since *w*Mel infections increase drastically as males age, a considerable shift in localization and density dynamics would be necessary. Microscopy assays will be required to test if *Wolbachia* and *cif* localization explains *w*Mel age-dependent CI-strength variation.

Second, age-dependent CI may be governed by developmental constraints of CI-susceptibility. For instance, the paternal grandmother age effect, where *Wolbachia*-infected sons of older virgin females cause stronger CI than sons of younger females, covaries with *Wolbachia* densities in embryos but not in adult males (30). Intriguingly, temperature-sensitive CI-strength variation in *Cardinium*-infected *Encarsia* wasps is also decoupled from symbiont densities, but CI strongly correlates with pupal development time (107, 108). *Cardinium* CI effectors likely have more time to interact with host targets at critical stages of pupal development when slowed by cool temperatures, despite lower *Cardinium* density (107, 108). These studies suggest that sperm are modified in spermatogenesis before adult eclosion, and that variation in symbiont densities during early development can contribute to CI-strength variation. If modified sperm are primarily produced during pupal or larval development, younger adult males would have a higher proportion of CI-modified sperm in their seminal vesicle than older males since older males continue to produce sperm as adults. Intriguingly, remating seems to weaken CI (86, 87), supporting this hypothesis. However, since CI strength decreases faster in *D. melanogaster* than in *D. simulans*, this hypothesis predicts that adult *D. simulans* sperm production is slower and/or CI modification occurs for an extended time. Functional work is necessary to determine if CI modification is developmentally restricted.

Finally, age-dependent CI may be related to the availability of CI-effector targets with male age and not the abundance of *cif* products. Indeed, the number of genes transcribed by *D. melanogaster* increases from 7,000 in embryos to over 12,000 in adult males, and nearly a third of genes are not expressed until the 3^rd^ larval instar (109). As adult males age, the number of transcribed genes continues to vary, though less so than during metamorphosis (109). These data support the possibility that host targets of CI may vary in abundance as males age. However, since transgenic *cif* expression can significantly enhance CI strength above wild-type levels (60), there are circumstances when natural *cif* expression is not high enough to saturate all targets. It is unknown if similar experimental approaches can strengthen age-dependent CI. More work will be necessary to determine the host genes that modify CI and how those factors vary in expression relative to CI strength.

### Age-dependent bacterial density covaries with immune expression, not phage or Octomom

We report a strong relationship between male age and *Wolbachia* densities that differ between systems: densities decrease in *w*Ri-infected *D. simulans* and increase in *w*Mel-infected *D. melanogaster*. Reports of age-dependent variation in *Wolbachia* densities across age in different tissues and sexes are common (51, 71, 90, 91, 110–112), but the basis of this variation remains unexplored. We investigated the cause(s) of this variation for the first time. We predicted that genes that covary with age-dependent densities might be causatively linked, although additional experiments will be necessary to confirm this. First, we tested whether phage or Octomom covary with age-dependent *Wolbachia* densities. Despite prior reports that phage WO of *Nasonia* and *Habrobracon Wolbachia* can regulate temperature-dependent *Wolbachia* densities (53, 66) and that Octomom copy number correlates with *w*MelCS and wMelPop densities (72, 73), we found that neither covaries with age-dependent *Wolbachia* densities in testes.

Next, we asked whether host immune gene expression correlates with age-dependent *Wolbachia* densities. We report that Relish expression, which activates antimicrobial peptide (AMP) production in the Imd pathway (101), strongly correlates with *w*Mel densities and is highest when *w*Mel densities are high. This result was surprising since we predicted that immune expression would hinder *Wolbachia* proliferation if it were correlated. Conversely, Relish does not vary with *D. simulans* male age and is only very weakly correlated with *w*Ri densities. It is plausible that the correlation between Relish expression and *w*Mel density represents a spurious and non-causative association. Additionally, Relish transcription does not necessarily equate to increased Relish activity and AMP production since endoproteolytic cleavage is necessary to activate the Relish protein (113), but future analysis of AMP expression will elucidate this. However, this correlation may represent a causative link between age-dependent *w*Mel densities and immune expression. We propose two non-exclusive hypotheses to explain this relationship.

First, *w*Mel rapidly proliferates as males age, elicit an immune response proportional to their infection density, but evade the effects of immune activation. *Wolbachia* synthesize lipid II (102–104), which is sufficient to activate the Imd pathway (104, 105), and increase AMP gene expression when transinfection into novel host backgrounds (114–117), suggesting that *Wolbachia* can trigger Imd activity. However, Relish and AMP expression do not vary with *Wolbachia* infection state (118–124) or density (84, 121) in natural *Wolbachia*-host associations. It has been proposed that *Wolbachia* evade the host immune system by residing in host-derived membranes or bacteriocyte-like cells (125, 126). Thus, the correlation between Relish expression and *w*Mel density may indicate that *w*Mel triggers the immune system but evades the immune response, preventing its densities from decreasing. Notably, since this hypothesis assumes that *w*Mel densities increase independent of Imd expression, it does not explain why *w*Mel densities increase with age or why age-dependent *w*Mel and *w*Ri densities differ.

Second, age-dependent Imd expression increases independent of *Wolbachia* but impacts *Wolbachia* densities. Aging in *D. melanogaster* is associated with increased expression of AMPs, Relish, and other immune genes (127–133). Counterintuitively, age also covaries with increased gut microbial loads and Imd activation in *D. melanogaster* (127–129, 134–136). Why gut bacterial loads increase with *D. melanogaster* age and immune expression remains unknown. However, age-dependent immune expression may damage the epithelium, lead to dysbiosis through differential effects on gut microbial members, alter gut tissue renewal and differentiation, and/or cause cellular inflammation (101, 137). In other words, the positive correlation between Relish expression and wMel density may be caused by off-target effects of immune expression on the cellular environment. To our knowledge, we report the first case where endosymbiont densities increase with age-dependent immune expression, suggesting that the cause(s) of age-dependent bacterial proliferation may apply to more than gut microbes. Functional assays, such as Relish knockdowns, will be necessary to causatively link male age-dependent *Wolbachia* densities and immune expression.

### Age-dependent CI strength could contribute to *Wolbachia* frequency variation in nature

We can consider our estimates of age-dependent CI strength in the context of an idealized discrete-generation model of *Wolbachia* frequency dynamics first proposed by Hoffmann et al. (1990). This model incorporates imperfect maternal transmission (*μ*), *Wolbachia* effects on host fitness (*F*), and the proportion of embryos that hatch in a CI cross relative to compatible crosses (*H*) (3). Across all experiments, CI strength (*s*_*h*_ = 1 - *H*) progressively decreases as males age (**Table S2**): *w*Mel CI strength decreases quickly (Day 0 *s*_*h*_ = 0.860; Day 8 *s*_*h*_ = -0.007) and *w*Ri CI strength decreases relatively slowly (Day 0 *s*_*h*_ = 0.991; Day 8 *s*_*h*_ = 0.244). Small negative values of *s*_*h*_ indicate that the CI cross has a slightly higher egg hatch than the compatible crosses.

*w*Ri occurs globally at high and relatively stable infection frequencies, consistent with generally strong CI (4, 26), while *w*Mel varies in frequency on several continents. In eastern Australia, *w*Mel frequencies range from ∼90% in the tropical north to ∼30% in the temperate south (34). While the factors that maintain this cline are unresolved, mathematical modeling suggests clinal differences in CI strength likely contribute (34). For example, CI must be essentially nonexistent (*s*_*h*_ << 0.05) to explain relatively low *w*Mel frequencies observed in temperate Australia, assuming little imperfect transmission (*μ =* 0.01 - 0.026) (138). Conversely, with *μ =* 0.026 and similarly low-to-nonexistent CI (*s*_*h*_ ≤ 0.055), large and positive *w*Mel effects on host fitness (*F* ∼ 1.3) are required to explain higher *w*Mel frequencies observed in the tropics. Explaining higher tropical frequencies becomes easier with stronger CI (*s*_*h*_ > 0.05) or more reliable *w*Mel maternal transmission (*μ <* 0.026) (Kriesner et al. 2016).

So what is *w*Mel CI strength in nature? Field-collected males from near the middle of the Australian cline to the northern tropics cause very weak (*s*_*h*_ ∼ 0.05) to no CI (Hoffmann et al. 1998). These, and other data from the middle of the cline (29), led Kriesner et al. (2016) to conjecture that the plausible range of *s*_*h*_ in subtropical/tropical Australian populations is *s*_*h*_ = 0 - 0.05, but < 0.1. In our study, only 6- (*s*_*h*_ = -0.006) and 8-day-old (*s*_*h*_ = -0.007) *w*Mel-infected males exhibited CI weaker than *s*_*h*_ = 0.1, suggesting that field-collected males causing little or no CI (138) are older than four days. Interactions among male age, temperature, remating, and other factors likely contribute to weaker CI in younger males (29, 37, 38, 53, 66, 86, 87). Future analyses to disentangle the contributions of male age and other factors to CI-strength variation are sorely needed. These estimates, along with estimates of *Wolbachia* transmission rate variation across genetic and abiotic contexts (22), are ultimately required to better understand *Wolbachia* frequency variation in host populations (7, 22, 24, 34, 139).

## Conclusions

Our results highlight that *Wolbachia* densities and *cif* expression from full-testes extracts are insufficient to explain age-dependent CI strength. While age-dependent CI strength in *w*Ri aligns with the bacterial density and CI gene expression hypotheses without the need to consider other factors, *w*Mel CI strength cannot be explained by either of these hypotheses. We propose that localization, development, and/or host genetic variation contribute to this relationship. Moreover, *w*Mel densities increase, and *w*Ri decrease as their respective hosts age. Neither phage WO nor Octomom explain age-dependent *Wolbachia* density, but variation in these systems covaries with the expression of the immune gene Relish. This represents the first report that the host immune system may contribute to variation in *Wolbachia* density in a natural *Wolbachia*-host association. This work motivates an extensive analysis of *Wolbachia* and *cif* expression in the context of localization and development and a thorough investigation of the relationship between host genes and *Wolbachia* density and CI phenotypes. Finally, Incorporating the age-dependency of CI into future modeling efforts may help improve our ability to explain temporally and spatially variable *Wolbachia* infection frequencies, as incorporating temperature effects on *w*Mel-like *Wolbachia* transmission has (22, 24, 140). Ultimately this will help explain *Wolbachia’s* status as the most prevalent endosymbiont in nature.

## Materials and Methods

### Fly lines

All fly lines used in this study are listed in **Table S4**. Uninfected flies were derived via tetracycline treatment in prior studies (16, 60). Tetracycline cleared lines were used in experiments over a year after treatment, mitigating the effects of antibiotics on mitochondria (141). We regularly confirmed infection status by using PCR to amplify the *Wolbachia* surface protein (*wsp*). An arthropod-specific 28S rDNA was amplified in a separate reaction and served as a control for DNA quality and PCR inhibitors (24, 142). The *y*^*1*^*w*^*1*^ *D. melanogaster* line was confirmed to be *w*Mel infected, as opposed to *w*MelCS, using IS5-WD1310 primers (143). DNA was extracted for infection checks using a squish buffer protocol. Briefly, flies were homogenized in 50 uL squish buffer per fly (100mL 1M Tris-HCL, 0.0372g EDTA, 0.1461g NaCl, 90 mL H_2_O, 150uL Proteinase K), incubated at 65°C for 45m, incubated at 94°C for 4m, centrifuged for 2m, and the supernatant was used immediately for PCR.

### Fly care and maintenance

Flies were reared in vials with 10mL of food made of cornmeal (32.6%), dry corn syrup (32%), malt extract (20.6%), inactive yeast (7.8%), soy flour (4.5%), and agar (2.6%). Fly stocks were maintained at 23°C between experiments. Flies used for virgin collections were reared at 25°C, virgin flies were stored at 25°C, and experiments were performed at 25°C. Flies were always kept on a 12:12 light:dark cycle. Flies were anesthetized using CO_2_ for virgin collections and dissections. During hatch-rate assays, flies were mouth aspirated between vials.

### Hatch-rate assays

CI manifests as embryonic death. We measured CI as the percentage of embryos that fail to hatch into larva. Flies used in hatch rates were derived from vials where flies were given ∼24hr to lay to control for rearing density (88). In the morning, virgin 6-8 day females were added individually to vials containing a small ice cream spoon filled with fly food. Spoon fly food was prepared as described above, but with blue food coloring added, 0.1g extra agar per 100mL of food, and fresh yeast smeared on top. After 4-5hr of acclimation, a single virgin male was added to each vial. The age of virgin males varied by experiment and cross. Paternal grandmother age was not controlled, but paternal grandmothers were non-virgin when setting up vials for fathers. Since *Wolbachia* densities associated with older paternal grandmothers are reduced upon mating (30), we do not expect variation in paternal grandmother *Wolbachia* densities across experiments or conditions. Vials with paired flies were incubated overnight at 25°C. Flies were then aspirated into new vials with a fresh spoon. Vials were incubated for another 24hr before flies were removed via aspirating. Embryos were counted on spoons immediately after flies were removed. After 48hr, the number of remaining unhatched eggs were counted. The percentage of embryos that hatched was then calculated.

### Relative abundance assays

Siblings from hatch-rate assays were collected for DNA extractions. Virgin males were anesthetized and testes were dissected in chilled phosphate-buffered saline (PBS). Five pairs of testes were placed into a single 1.5mL Eppendorf tube and stored at -80°C until processing. All tissue was collected the day after the hatch-rate setup. Tissue was homogenized using a pestle, and the DNeasy Blood and Tissue kit (Qiagen) was used to extract and purify DNA.

qPCR was used to measure the relative abundance of host, *Wolbachia*, phage WO, and Octomom DNA. Samples were tested in triplicate using Powerup SYBR Green Master Mix (Applied Biosystems), which contains a ROX passive reference dye. Primers were designed using Primer3 v2.3.7 in Geneious Prime (144). Host primers target an ultraconserved element (UCE) *Mid1* identified previously (96). Phage genes were also identified from prior works (100). Primers for *w*Mel’s phages target both WOMelA (WD0288) and WOMelB (WD0634), while those for *w*Ri are unique to a single phage haplotype. WORiA, WORiB, and WORiC were measured with *w*Ri_012460, *w*Ri_005590/*w*Ri_010250, and *w*Ri_006880 primers, respectively. Only *w*Mel has all eight Octomom genes (WD0507-WD0514) (71). We measured *w*Mel Octomom copy number using primers targeting WD0509. Primer sequences and PCR conditions are listed in **Table S5**. Fold difference was calculated as 2^-ΔΔCt^ for each comparison. A random sample in the youngest age group was selected as the reference.

### Gene expression assays

Siblings from hatch-rate assays were collected for RNA extractions. Virgin males were anesthetized, and testes were dissected in chilled RNase-free PBS. Fifteen pairs of testes were placed into a single 2mL tube with 200uL of Trizol and four 3mm glass beads. Tissue was kept on ice between dissections. Samples were then homogenized using a TissueLyser II (Qiagen) at 25Hz for 2m, centrifuged, and stored at -80°C until processing. All tissue was collected the day after the hatch-rate setup.

Samples were thawed, 200uL of additional Trizol was added, and tissue was further homogenized using a TissueLyser II at 25Hz for 2m. RNA was extracted using the Direct-Zol RNA Miniprep kit (Zymo Research) following the manufacturer’s recommendations, but with an extra wash step. On-column DNase treatment was not performed. Instead, the ‘rigorous’ treatment protocol from the DNA-free kit (Ambion) was used to degrade DNA in RNA samples. Samples were confirmed DNA-free using PCR and gel electrophoresis for an arthropod-specific 28S rDNA (24, 142). The Qubit RNA HS Assay Kit (Invitrogen) was used to measure RNA concentration. Samples within an experiment were diluted to the same concentration. RNA was converted to cDNA using SuperScript IV VILO Master Mix (Invitrogen) with either 200ng or 500ng of total RNA per reaction depending on the experiment. qRT-PCR was performed using 1ng of cDNA per reaction using Powerup SYBR Green Master Mix (Applied Biosystems). All samples were tested in triplicate.

Primers for expression included host reference, *Wolbachia* reference, *cif*, and host immune genes. Primers to *Drosophila* genes for qRT-PCR were selected from FlyPrimerBank (145). Since *Drosophila* expression patterns change with age (109), a host gene that is invariable with male age was selected to act as a reference gene for relative expression analyses. We selected an invariable gene using the *Drosophila* Gene Expression Tool (DGET) to retrieve modENCODE gene expression data for ribosome and cytoskeletal genes (146). DGET reports expression as Reads Per Kilobase of transcript, per million mapped reads (RPKM), and included data for adult males 1, 5, and 30 days after eclosion. β-spec (1 Day = 81 RPKM, 5 Day = 80, 30 Day = 79) was selected because it is largely invariable across age. Our results confirm invariable expression across male age (**Fig. S2E**; **Fig. S3L**). *D. melanogaster* and *D. simulans* are identical across *βspec* primer binding sequences. All other primers were designed using Primer3 in Geneious Prime (144) and are listed in **Table S5**. Fold difference was calculated as 2^-ΔΔCt^ for each comparison. A random sample in the youngest age group was selected as the reference.

### Statistical analyses

All statistics were performed in R (147). Hatch-rate, relative-abundance, and expression assays were analyzed using a Kruskal-Wallis followed by a Dunn’s test with corrections for multiple comparisons. Kruskal-Wallis and Dunn’s *P*-values are reported in **Table S1**. Correlations between hatch rate and expression or relative abundance measures were performed using Pearson and Spearman correlations in GGPubR (148). Correlation statistics are reported in **Table S3**. 95% confidence intervals were calculated using the classic MeanCI function in DescTools (149). 95% BCa intervals were calculated using boot.ci in boot (150). Samples with fewer than ten embryos laid were excluded from hatch-rate analyses. Samples with a C_q_ standard deviation exceeding 0.4 between triplicate measures were excluded from qPCR and qRT-PCR analyses. Figures were created using GGPlot2 (151), and figure aesthetics were edited in Affinity Designer 1.8 (Serif Europe, Nottingham, UK).

## Data availability

All data are made publicly available in the supplement of this manuscript.

## Acknowledgments

We thank Michael Turelli for helpful feedback on our experimental design. We also thank Will Conner for help identifying phage gene targets for qPCR, Mike Hague for support with BCa estimates of *H*, Kelley Van Vaerenberghe and John Statz for review of earlier versions of the manuscript, and Tim Wheeler for assisting in the laboratory. Seth Bordenstein provided comments on a preprint. This work was supported by a National Institutes of Health R35 GM124701 to BSC and a National Science Foundation Postdoctoral Research Fellowship DBI-2010210 to JDS. Any opinions, findings, conclusions, or recommendations expressed in this material are those of the authors(s) and do not necessarily reflect the views of the National Institutes of Health or the National Science Foundation.

## Supporting information

**Figure S1. Testing the bacterial density and Octomom copy number hypotheses for CI strength variation in young *w*Mel-infected *D. melanogaster***. Fold change across male age for (A) *w*Mel *ftsZ* relative to *D. melanogaster* UCE and (B) Octomom gene WD0509 to *w*Mel *ftsZ*. Letters above data represent statistically significant differences based on α=0.05 calculated by Dunn’s test with correction for multiple comparisons between all groups —crosses that do not share a letter are significantly different. Fold change was calculated as 2^-ΔΔcq^. *P*-values are reported in Table S1.

**Figure S2. Testing the *cif*-expression hypothesis for *w*Mel CI strength variation**. Fold change across male age for the relative expression of (A) *cifB*_*wMel[T1]*_ to *D. melanogaster βspec* and (B) *cifB*_*wMel[T1]*_ to *w*Mel *ftsZ*. Raw C_q_ values for (C) *cifA*_*wMel[T1]*_, (D) *cifB*_*wMel[T1]*_, (E) *D. melanogaster βspec*, and (D) *w*Mel *ftsZ*. Letters above data represent statistically significant differences based on α=0.05 calculated by Dunn’s test with correction for multiple comparisons between all groups —crosses that do not share a letter are significantly different. Fold change was calculated as 2^-ΔΔcq^. *P*-values are reported in Table S1.

**Figure S3. Testing the *cif*-expression hypothesis for *w*Ri CI strength variation**. Fold change across male age for the relative expression of (A) *cifB*_*wRi[T1]*_ to *D. simulans βspec*, (B) *cifB*_*wRi[T1]*_ to *w*Ri *ftsZ*, (C) *cifA*_*wRi[T2]*_ to *D. simulans βspec*, (D) *cifA*_*wRi[T2]*_ to *w*Ri *ftsZ*, (E) *cifB*_*wRi[T2]*_ to *D. simulans βspec*, (F) *cifB*_*wRi[T2]*_ to *w*Ri *ftsZ*, and (G) *cifA*_*wRi[T1]*_ to *cifA*_*wRi[T2]*_. Raw C_q_ values for (H) *cifA*_*wRi[T1]*_, (I) *cifB*_*wRi[T1]*_, (J) *cifA*_*wRi[T2]*_, (K) *cifB*_*wRi[T2]*_, (L) *D. simulans βspec*, and (M) *w*Ri *ftsZ*. Letters above data represent statistically significant differences based on α=0.05 calculated by Dunn’s test with correction for multiple comparisons between all groups —crosses that do not share a letter are significantly different. Fold change was calculated as 2^-ΔΔcq^. *P*-values are reported in Table S1.

**Figure S4. Testing the phage density model for *Wolbachia*-density variation**. Fold change across male age for the relative abundance of (A) WOMelA/B to *D. melanogaster* UCE in the 0-8 day old male age experiment, (B) WOMelA/B to *D. melanogaster* UCE in the 0-3 day old male age experiment, (C) WOMelA/B to *w*Mel *ftsZ* in the 0-3 day old male age experiment, (D) WORiB to *D. simulans* UCE, (E) WORiA to *w*Ri *ftsZ*, and (F) WORiC to *w*Ri *ftsZ*. Letters above data represent statistically significant differences based on α=0.05 calculated by Dunn’s test with correction for multiple comparisons between all groups —crosses that do not share a letter are significantly different. Fold change was calculated as 2^-ΔΔcq^. *P*-values are reported in Table S1.

**Table S1. *P*-values associated with all statistical comparisons**.

**Table S2. Parameter estimates of H and Sh**. Hatch-rate data used for estimates are derived from experiments in Figure 2.

**Table S3. Correlations between relative abundance or relative expression and hatch rate**. Correlations are organized by the relevant hypothesis: bacterial density, phage density, host immunity, Octomom, and *cif* expression. Factors positively correlated with hatch rate are negatively correlated with CI strength, and vice versa. Pearson and Spearman *P*-values are bold if < 0.05, underlined if < 0.01, and double-underlined if < 0.001. *w*Mel “old cohort” refers to the experiment presented in Fig. 2B where 0-, 2-, 4-, 6-, and 8-day-old males are tested for CI. *w*Mel “young cohort” refers to the experiment in Fig. 2A where 0-, 1-, 2-, and 3-day-old males are tested for CI. *w*Ri data comes from siblings of males used in Fig. 2C. Median hatch rates were used for correlations and were taken from relevant experiments in Fig. 2.

**Table S4. *Drosophila* lines used in this study. Table S5. Primers used in this study**.

**Supporting data file 1. All hatch-rate and qPCR data generated in this study**.

