## Supplementary figures and images for "Male age and *Wolbachia* dynamics: Investigating how fast and why bacterial densities and cytoplasmic incompatibility strengths vary"

### Figures S1-S4

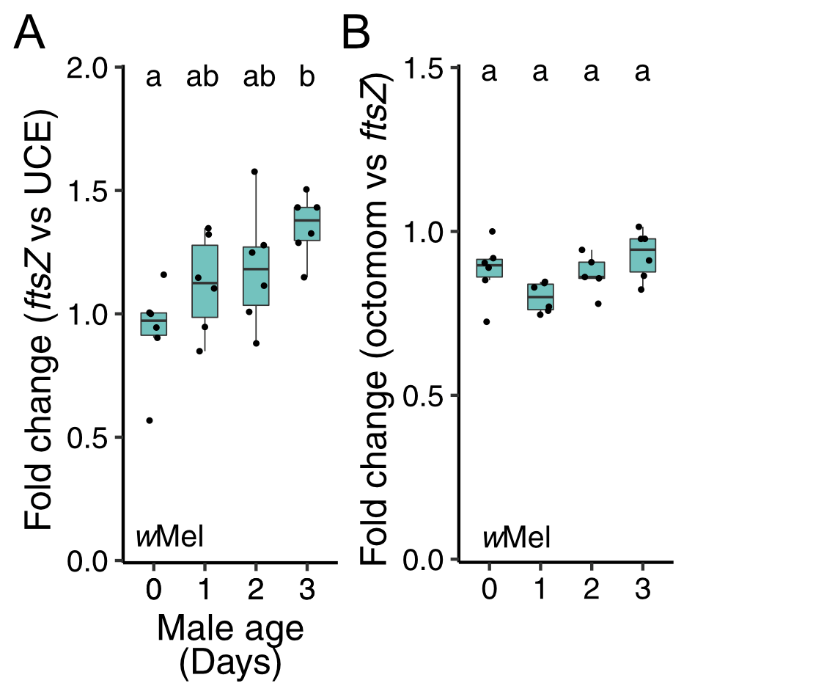

Figure S1.

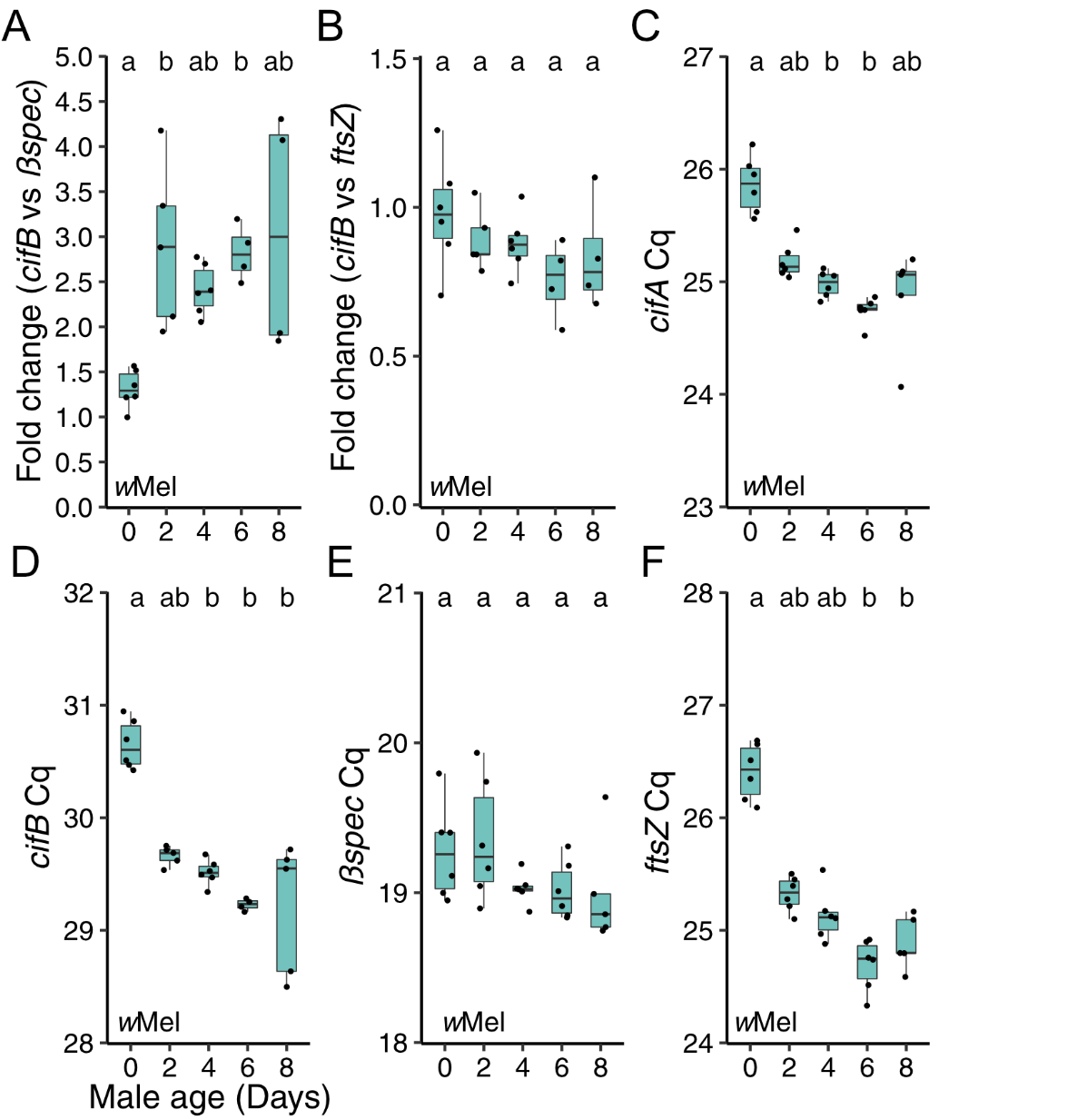

Figure S2.

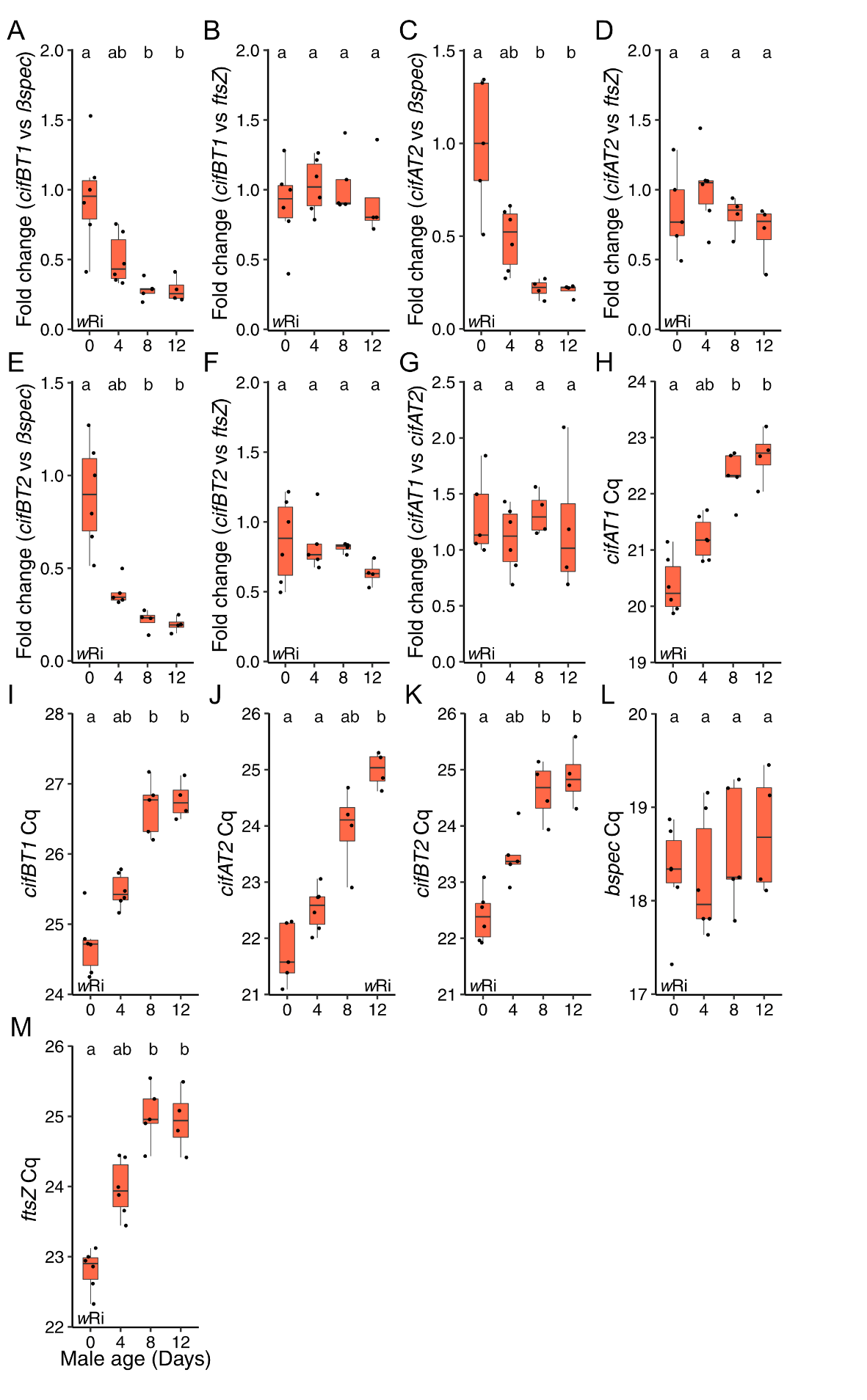

Figure S3.

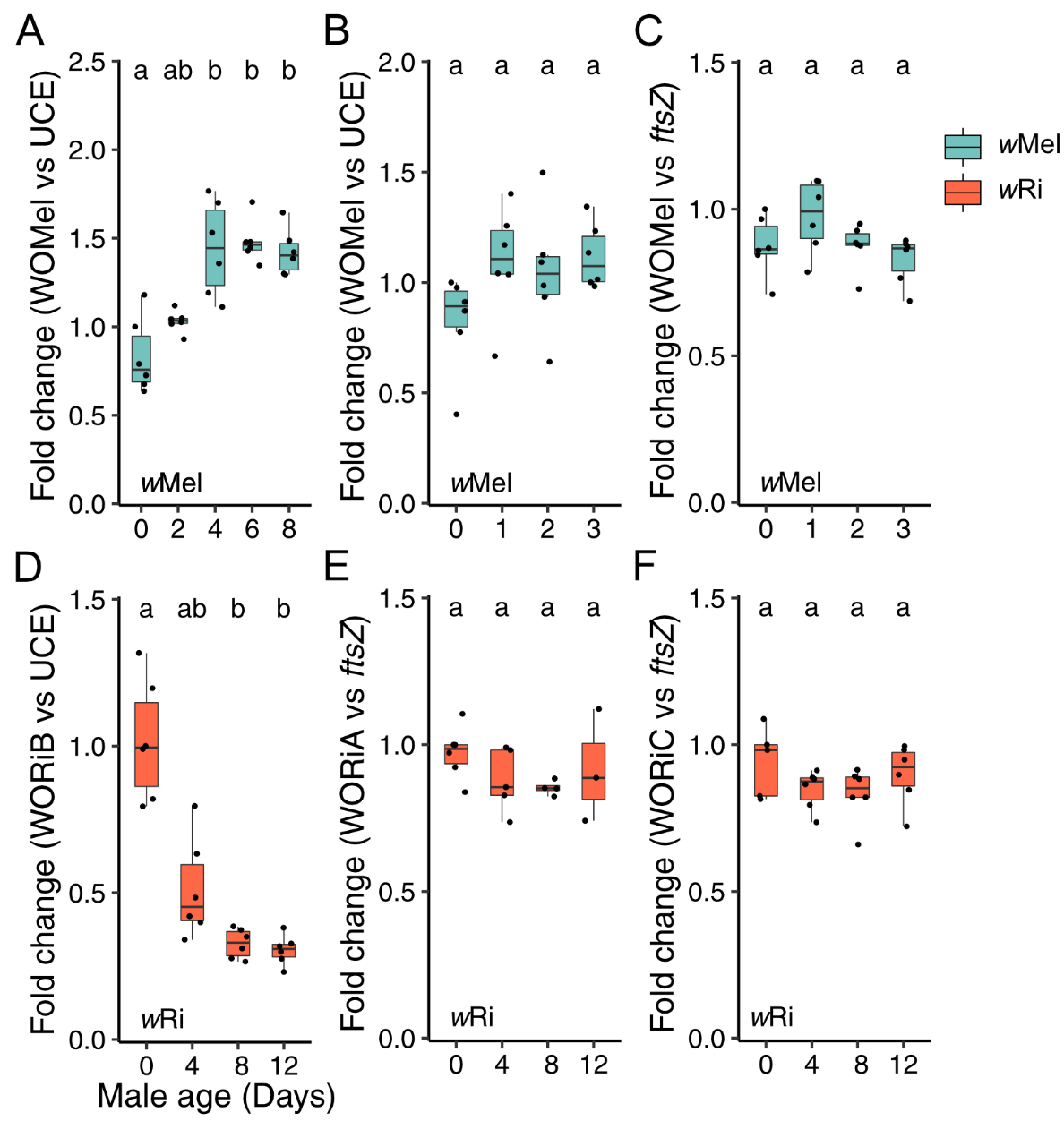

Figure S4.
